# Vagus nerve stimulation rescues persistent pain following orthopedic surgery in adult mice

**DOI:** 10.1101/2023.05.16.540949

**Authors:** Pau Yen Wu, Ana Isabel Caceres, Jiegen Chen, Jamie Sokoloff, Mingjian Huang, Gurpreet Singh Baht, Andrea G Nackley, Sven-Eric Jordt, Niccolò Terrando

## Abstract

Postoperative pain is a major clinical problem imposing a significant burden on our patients and society. Up to 57% of patients experience persistent postoperative pain 2 years after orthopedic surgery [49]. Although many studies have contributed to the neurobiological foundation of surgery-induced pain sensitization, we still lack safe and effective therapies to prevent the onset of persistent postoperative pain. We have established a clinically relevant orthopedic trauma model in mice that recapitulates common insults associated with surgery and ensuing complications. Using this model, we have started to characterize how induction of pain signaling contributes to neuropeptides changes in dorsal root ganglia (DRG) and sustained neuroinflammation in the spinal cord [62]. Here we have extended the characterization of pain behaviors for >3 months after surgery, describing a persistent deficit in mechanical allodynia in both male and female C57BL/6J mice after surgery. Notably, we have applied a novel minimally invasive bioelectronic approach to percutaneously stimulate the vagus nerve (termed pVNS) [24] and tested its anti-nociceptive effects in this model. Our results show that surgery induced a strong bilateral hind-paw allodynia with a slight decrease in motor coordination. However, treatment with pVNS for 30-minutes at10 Hz weekly for 3 weeks prevented pain behavior compared to naïve controls. pVNS also improved locomotor coordination and bone healing compared to surgery without treatment. In the DRGs, we observed that vagal stimulation fully rescued activation of GFAP positive satellite cells but did not affect microglial activation. Overall, these data provide novel evidence for the use of pVNS to prevent postoperative pain and may inform translational studies to test anti-nociceptive effects in the clinic.

## 1. Introduction

Pain is a critical component following orthopedic surgery, and its chronification can significantly impact recovery and quality of life. Severe postoperative pain frequently develops into chronic pain and is common after orthopedic procedures [3]. More than 5 million Americans across all age groups undergo orthopedic surgery every year to repair common bone injuries, such as fractures, and to replace joints [35; 36]. Poorly managed postoperative pain interferes with successful rehabilitation and optimal healing, leading to long-lasting loss of bone and muscle mass, and eventually chronic pain [36; 49]. Current treatment strategies for orthopedic postsurgical pain employ NSAIDs and opioids that carry the risks for adverse side effects, including gastrointestinal, cardiovascular, hepatic, bone healing as well as addiction [41]. Here, we applied a model of a commonly performed orthopedic surgical procedure to study the impact of trauma on pain behavior, neuroinflammation, motor coordination, and bone healing and tested a bioelectronic approach as a novel rescuing strategy.

Bioelectronic medicine, including approaches that leverage electrical stimulation of the vagus nerve (VNS) are commonly used to treat conditions such as epilepsy and depression [32]. VNS has well-established anti-inflammatory effects through regulation of neuro-immune signaling and activation of the cholinergic anti-inflammatory pathway [21]. This inflammatory reflex plays a crucial role in monitoring peripheral inflammatory responses [44]. Notably, cholinergic stimulation modulates the immune response in several conditions ranging from sepsis to rheumatoid arthritis [57], and the implications for perioperative care have been recently highlighted [52]. Currently, no study has evaluated the effectiveness of VNS on preventing chronic pain and modulating the activity of immune cells in pain-sensing primary afferent neurons. We have implemented a minimally invasive yet targeted approach to stimulate the vagus perioperatively (pVNS) and demonstrated engagement of the cholinergic anti-inflammatory pathway as well as modulation of neuroinflammation and cognitive outcomes[24].

Glia cells exerts important roles and the functions of DRG macrophages and satellite glia in pain processing are well-appreciated [29]. Microglia in the spinal cord have a crucial role in the development of neuropathic pain following nerve injury. Upregulation of markers such as Iba1 and CD11b after nerve injury have been often observed in the spinal cord [11]. In addition, neuromodulators such as ATP, colony-stimulating factor–1 (CSF1) and chemokines like C-C motif ligand 2 / monocyte chemoattractant protein 1 (CCL2/MCP-1), Brain derived neurotrophic factor (BDNF), tumor necrosis factor (TNF) and interleukin 1β (IL-1β) have been all involved with microglial-meditated pain chronification [8; 15; 18; 19]. Astrocytes have also been shown to interact with nociceptive neurons to amplify persistent pain signaling. Signal molecules such as ATP and CCL2 released form the injury site can trigger astrocytes to initiate a vicious cycle and further activate nociceptive neurons *via* glutamate, CCL2, CXCl1 or TSP4 [28; 30]. Currently, limited research focuses on the long-term response of glial cells in the DRG, especially satellite glial cells. Here, we have applied a novel, minimally invasive pVNS approach to ascertain its putative analgesic effects following orthopedic surgery and measured neuroinflammatory changes in the DRG of male and female adult mice at the point when pVNS improved pain behavior.

## 2. Methods

### 2.1 Animals

C57BL6/J male and female mice 12 weeks old were obtained from The Jackson Laboratory (Bar Harbor, ME). Animals were housed 3–5 per cage under rolled temperature and humidity with a 14:10 h light:dark cycle. Mice were fed standard rodent chow (Prolab RMH3500, Autoclavable; LabDiet, St. Louis, MO) and had *ad libitum* access to food and water. All experiments were conducted under an approved protocol from the Institutional Animal Care and Use Committee at Duke University Medical Center and under the guidelines described in the National Science Foundation “Guide for the Care and Use of Laboratory Animals” (2011).

### 2.2 Fos-TRAP2; Ai14 and 4-OHT administration

FosTRAP (Fos the targeted recombination in active populations) mice were a gift of Dr. Fan Wang (MIT) and used as recently reported [9; 10]. The FosTRAP mice were kept in single cages with at least two environmental enrichments for a minimum of 10 days prior the day of 4-Hydroxytamoxifen (4-OHT) injection. The 4-OHT (4-OHT; Sigma, MO, USA Cat# H6278) was dissolved in ethanol to create a 20mg/ml stock solution, which was then divided and frozen in – 20°C. On the day of administration, the 4-OHT was added to a sunflower seed oil solution (Sigma, MO, USA Cat # S5007) and heated to 50°C to remove the ethanol, resulting in a final 4-OHT solution with a concentration of 10 mg/ml. Any unused solution was discarded at the end of the day. The mice received an intraperitoneal 4-OHT injection 30 minutes before undergoing 30 minutes of pVNS treatment or sham. Both pVNS and sham group received sevoflurane (UPS, Covetrus North America, OH, USA) anesthesia. The mice were returned to the same single-housing cage for 10 days to minimize any sources of distress before termination.

### 2.3 Orthopedic surgery

Tibial surgery was performed as described [59]. Mice were anesthetized with isoflurane via a low-flow digital anesthesia system (SomnoSuite apparatus, Kent Scientific Corporation, Torrington, CT, USA). Body temperature was maintained at 36.5 ± 0.6°C using a homoeothermic pad system (Kent Scientific Corporation). Muscles were dissociated following an incision on the left hind paw. A 0.38-mm stainless steel pin was inserted into the tibia intramedullary canal, followed by osteotomy, and the incision was closed with 6-0 Prolene suture. Control mice used in this study were home-cage naïve mice. In the persistence study (Fig 1), the sham mice was were anesthetized with isoflurane and body temperature was maintained at 36.5 ± 0.6°C using a homoeothermic pad system (Kent Scientific Corporation). An incision was made on the skin of the mice to expose the tibial muscle, after which the skin was stitched back together with 6-0 Prolene suture.

**Fig 1.**
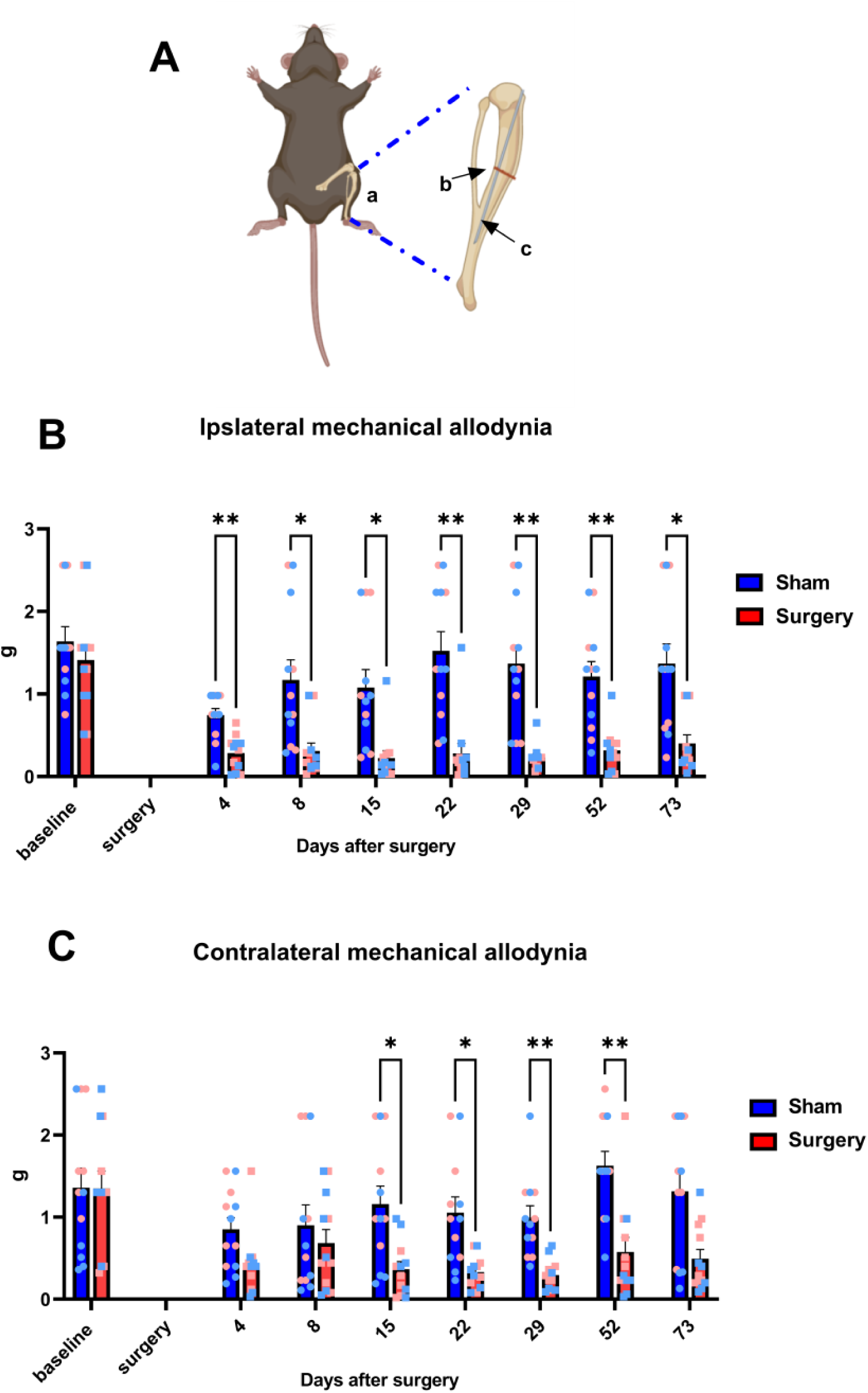
Orthopedic surgery induces persistent mechanical allodynia. A) Figure of tibia fracture with supported pin in bone marrow as surgery model. Both ipsilateral (B) and contralateral (C) hind-paw surgery induced persistence mechanical allodynia for 73 and 52 days respectively. Analyzed by Two-way ANOVA with Bonferroni posttest. *, **p < 0.05, 0.01 surgery vs. Sham surgery. Both sex was included in each group. N=5-6 in each sex/group.

### 2.4 Percutaneous vagus nerve stimulation (pVNS)

pVNS was performed as pervious described [24]. Mice were under anesthesia with 6% induction sevoflurane (Covertrus, Dublin, OH, USA) then were placed on a heatpad (39 °C) with nose-cone delivered 3-5% sevoflurane to maintain general anesthesia. Pulse-oximetry (MouseSTAT Jr.; Kent Scientific Corporation, Torrington, CT) was applied to monitor the real time heart rate. VNS approach under ultrasound guidance and non-invasive pulse oximetry monitoring ascertain the effects of vagus stimulation on sympathetic activity (0.2-1mA to achieve 90% bradycardia threshold, 20 Hz, 0.3 ms/pulse biphasic).

### 2.5 Hind paw mechanical withdrawal threshold

The 50% paw withdrawal threshold was tested before surgery and VNS treatment and on days 4, 8, 15, 22, 29 post-surgery. There were two acclimation sessions on 3 and 5 days before the baseline hindpaw mechanical withdrawal threshold testing. During the acclimation session, mice were place in the testing room for half hour in home cage and another one hour on testing mesh wire table. On the day of testing, mice were acclimated in home cage for 30 mins then placed in individual clear plastic chambers (11 × 5 × 3.5cm) on a 55cm-high wire mesh table and allowed to acclimate for 30 mins. The up-down method was performed using a series of monofilaments (1.65, 2.36, 2.83, 3.22, 3.61, 4.08, 4.31, 4.74 g; North Coast Medical, Inc Morgan Hill, CA, USA) applied to the left or right hind paw. The test started with application of the 3.22g monofilament. A negative response was followed by the next larger monofilament, whereas a positive response (brisk withdrawal of the paw) was followed by the next smaller monofilament. Four additional filaments were tested after the first positive response. The 50% withdraw threshold was calculated for each mouse and group means were determined as previously described [5].

### 2.6 Rotarod performance test

On 18, 25 and 32 days after orthopedic surgery, mice were placed on the rotarod machine (LE8500, Panlab S.L.U., Barcelona, Spain) three times (mix 60 seconds) at speed of 4 rpm for acclimatization 2-3 hours before the test. During the test mice were place on rotarod machine, starting with 4 rpm with 2 ACC setting. The time spent on the rod for each mouse was recorded. Each mouse was tested for three trials with more than 5 minutes resting time between each trial. The average of each trial indicated the gross motor performance of mice.

### 2.7 Tissue Collection

Animals were euthanized under deep isoflurane anesthesia via trans-cardiac perfusion with ice-cold phosphate buffered saline (PBS; 0.1 M, pH 7.4; 50 ml), followed by ice-cold paraformaldehyde (4% PFA in PBS; 50 ml). Brains and DRGs were then dissected, placed in 4% PFA overnight, washed with PBS, placed in 30% sucrose solution for 3 days, and then stored in OCT at −80 °C until sectioning. Post-fixed brains (PFA 4%) were coronally sectioned at either 30 μm (IHC) with a cryostat (Leica CM 1950, Leica Biosystems, Deer Park, IL, USA). Mice left tibia was dissected and placed in 4% PFA then follow 2.10 Bone histology analysis method.

### 2.8 Immunohistochemistry

Free floating sections were washed 3 times with PBS (15 min each), incubated in 0.3% triton in PBS for 20 minutes, blocked for 1 hour in blocking buffer solution (2.5% BSA; 0.3% Triton), and incubated overnight at 4°C with the primary antibody diluted in the blocking buffer solution. The next day, sections were washed 3 times with the blocking buffer solution (15 min each) and subsequently incubated for 1h at RT with the secondary antibody diluted in the blocking buffer solution: Donkey anti Rabbit 647 (1:1000, AB150075, ABCAM, Cambridge, UK); Donkey anti Mouse 594, anti Goat 488, anti Rabbit 488 (1:1000, A21203, A11055, A21206, Invitrogen, Thermo Fischer Scientific). Lastly, sections were washed 2 times with PBS (10 mins each) and finally mounted a slide covered with DAPI mounting media (#F6057, Sigma-Aldrich).

### 2.9 Image analysis

Image stacks of the regions of interest were acquired with a 20x, 40x or 63x objective using the Zeiss inverted 880 confocal microscope. Maximum intensity projections were applied by Zeiss black software follow by ImageJ’s threshold analysis and background subtraction. Automatic algorithms were determined to match best the cell morphology and decrease the noise/signal ratio as described previously [20].

### 2.10 Bone histology analysis

Fracture calluses were dissected and fixed in 10% Zn-formalin at room temperature for 5 days. MicroCT analysis was conducted using a Scanco vivaCT 80 (Scanco Medical, Brüttisellen, Switzerland) at a scan resolution of 8µm. Fractured tibiae were scanned, and the midpoint of the fracture callus was identified. Calluses were analyzed 1mm proximal and 1mm distal from the fracture site and assessed for total volume (TV) and bone volume (BV) in mm3, percent bone volume per total volume (%BV/TV), and tissue mineral density (TMD) in mg HA/mm3. Fixed fracture calluses were decalcified using 12% EDTA pH 7.4, cleared of EDTA, and embedded into paraffin. Sections were cut at a thickness of 5µm and stained using Alcian Blue/Hematoxylin/Orange-G to visualize bone and cartilage. A minimum of five sections were used to conduct computer-assisted histomorphometry analysis, and results were presented as an amount relative to the total area of the fracture callus, as previously described [23].

### 2.11 Statistics

Mouse data were analyzed with one-way or two-way analysis of variance (ANOVA), with or without repeated measures, followed by Tukey’s or Bonferroni’s posttest, as denoted in the figure legends. Bone and NTS Td-tomato expression data was analyzed by t-test. All hypothesis testing was 2-tailed and statistical significance was set at alpha = 0.05. Mouse data were analyzed using GraphPad Prism (GraphPad v9.4.1; GraphPad Software, San Diego, CA, USA). All data were expressed as mean ± SEM.

## Results

### 3.1 Persistent allodynia following orthopedic surgery

We used a well-established model of orthopedic surgery with intramedullary fixation of the tibia to longitudinally evaluate pain behavior in adult male and female mice (Fig. 1A). We observed persistent allodynia both in the ipsilateral and contralateral paw up to 73 days and 52 days, respectively. For the ipsilateral hind paw a significantly higher (2way-Anova with Bonferroni posttest, F (1, 22) = 27.16, p <0.0001) withdrawal threshold was observed following surgery compared to shams on Day 4 (p=0.0013, MD=0.46, 95% CI [0.1559 to 0.7641]), Day 8 (p=0.0422, MD=0.8592, 95% CI [0.02225 to 1.696]), Day 15 (p=0.0206, MD=0.855, 95% CI [0.1033 to 1.607]), Day 22 (p=0.0016, MD=1.243, 95% CI [0.4241 to 2.063]), Day 29 (p=0.0028, MD=1.134, 95% CI [0.3745 to 1.894]), Day 52 (p=0.0036, MD=0.8933, 95% CI [0.2616 to 1.525]) and Day 73 (p=0.0162, MD=0.9667, 95% CI [0.1425 to 1.791]) (Fig. 1B). For the contralateral hind paw allodynia started at a later time point, day 15 (p=0.0397, MD=0.7925, 95% CI [0.02713 to 1.558]), compared to the ipsilateral side, lasting up to day 52 (p=0.0029, MD=1.05, 95% CI [0.2964 to 1.804]) (Fig. 1C). Overall, these data indicated bilateral persistent allodynia following orthopedic surgery in adult mice.

### 3.2 Tracing neuronal activation after pVNS

We applied a minimally invasive pVNS approach under ultrasound guidance and non-invasive pulse oximetry monitoring to effectively and reliably stimulate the vagus nerve in anesthetized mice. During the stimulation, a bipolar electronic needle was placed on right vagus nerve guided by mice neck ultrasound while mice were immobile under sevoflurane anesthesia (Fig. 2A,B). Non-invasive pulse oximetry was used to monitor mice heart rate and bradycardia to confirm the effectiveness of vagus nerve stimulation (Fig. 2C). We confirmed activation of vagal nerve afferent signaling to the Nucleus Tractus Solitarius (NTS) following pVNS (0.2-1mA to achieve 90% bradycardia threshold, 20 Hz, 0.3 ms/pulse biphasic) in FosTRAP mice as demonstrated by a robust induction of Fos+ neurons labeled by Td-tomato (Unpaired t test, p= 0.011, 95% CI 2.245 to 10.76) (Fig 2 D,E,F). Overall, FosTRAP mice provided further validation for the activation of the NTS following pVNS exposure.

**Fig 2.**
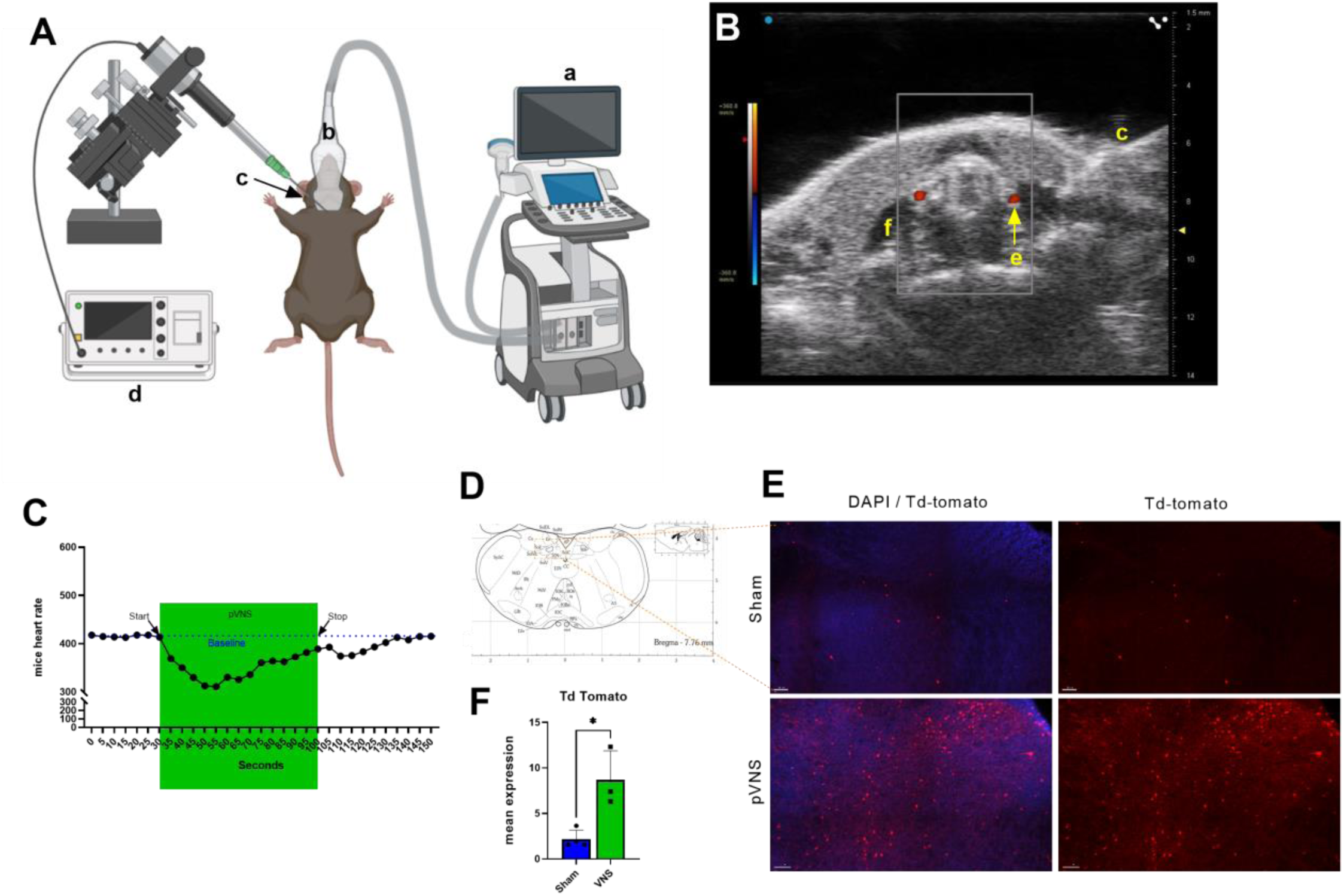
pVNS setup and ultrasound imaging. A) pVNS setup with a: ultrasound image in the monitor. b: ultrasound probe placed on mice neck. c: electronic needle. d: electronic stimulator. B) Doppler ultrasound imaging. c: electronic needle. e: carotid artery. f: cervical muscles. C) pVNS induced bradycardia on mice. D) NTS reconstructed from Paxinos and Franklin’s the Mouse Brain in Stereotaxic Coordinates. E) pVNS increase td-tomato expression on FosTrap2 mice in nucleus of solitary tract. F) Quantification data from panel E.

### 3.3 Repetitive pVNS improve surgery-induced allodynia without impairing healing

Next, we investigated the effects of 30 minutes weekly pVNS on postoperative pain. On the injury day, one session of 30 min pVNS treatment was applied just after surgery. Mice were randomly assigned to three groups as described in Fig. 3A. Three weeks after surgery, on Day 22 (p=0.0014, MD=−1.402, 95% CI [−2.253 to −0.5516]) and Day 29 (p=<0.0001, MD=−2.131, 95% CI [−3.052 to −1.209]), we started to observe a separation between pVNS-treated and control mice after surgery. (Fig. 3A, B). Notably, we did not observe sex difference in response to pVNS in either ipsilateral (2way-Anova, surgery p=0.9636 95%CI= −0.03952 to 0.04127, Surgery+VNS p= 0.5543 95%CI= −0.3652 to 0.6527) and contralateral sides (2way-Anova, surgery p=0.7085 95%CI= −0.1662 to 0.2382, Surgery+VNS p= 0.2633 95%CI= −0.2603 to 0.8798) (Fig. 3C, D). Overall, we concluded that pVNS is effective in reducing postoperative pain in both male and female mice.

**Fig 3.**
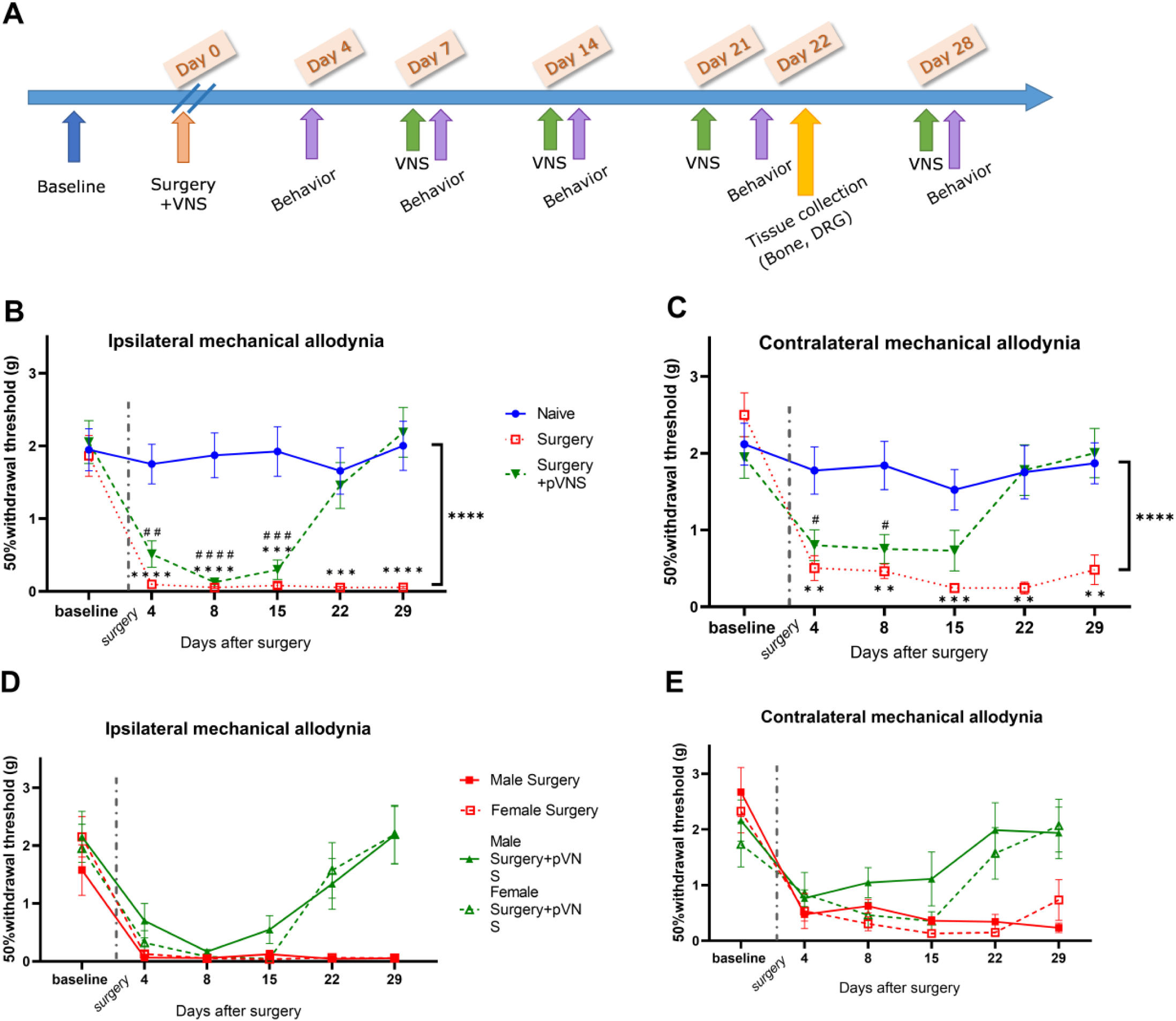
pVNS rescued surgery-induced bilateral mechanical allodynia. A) Surgery, pVNS, behavior experiment and tissue collection timeline. B,C) VNS reversed the surgery-induced chronic mechanical hind-paw allodynia after three weeks in both ipsilateral (B) and contralateral paw(C). There is no significant sex different between Male surgery group and female surgery groups or male surgery+VNS and female surgery+VNS groups in both ipsilateral (D) and contralateral (E) hindpaw. Analyzed by Two-way ANOVA with Bonferroni posttest. **, ***, **** p < 0.01, 0.001, 0.0001 Surgery vs. Naive; #, ##, ###, #### p < 0.05, 0.01, 0.001, 0.0001 Surgery+VNS vs. Naïve, both sex was included with N=6-8 in each sex/group.

### 3.4 Multiple pVNS treatments escalate bone healing and improve motor correlation function

We then examined whether multiple pVNS treatments were safe in respect to bone healing after orthopedic surgery. pVNS did not have a significant impact on the total volume of the callus nor on the bone volume (Fig 4. B, D). Interestingly, pVNS increased bone ratio (BV/TV) compared to surgery only mice (Unpaired t test, p= 0.0036, 95% CI 0.04039 to 0.1776) (Fig 4. D). Both sexes demonstrated no significant difference in rotarod testing. In fact, the data suggest improve rotarod performance on surgery+VNS group compared to surgery only group on postoperative day 10 (p=0.0328, MD=6.417, 95% CI [12.26 to 0.5779]), day 18 (p=0.0262, MD=7.25, 95% CI [13.71 to 0.7866]) and day 32 (p=0.0087, MD=6.292, 95% CI [11.07 to 1.516]) (Fig. 4 F,G,H). Overall, this indicates that pVNS does not negatively impact bone healing and repetitive stimulation is safe in preventing conversion to chronic pain.

**Fig 4.**
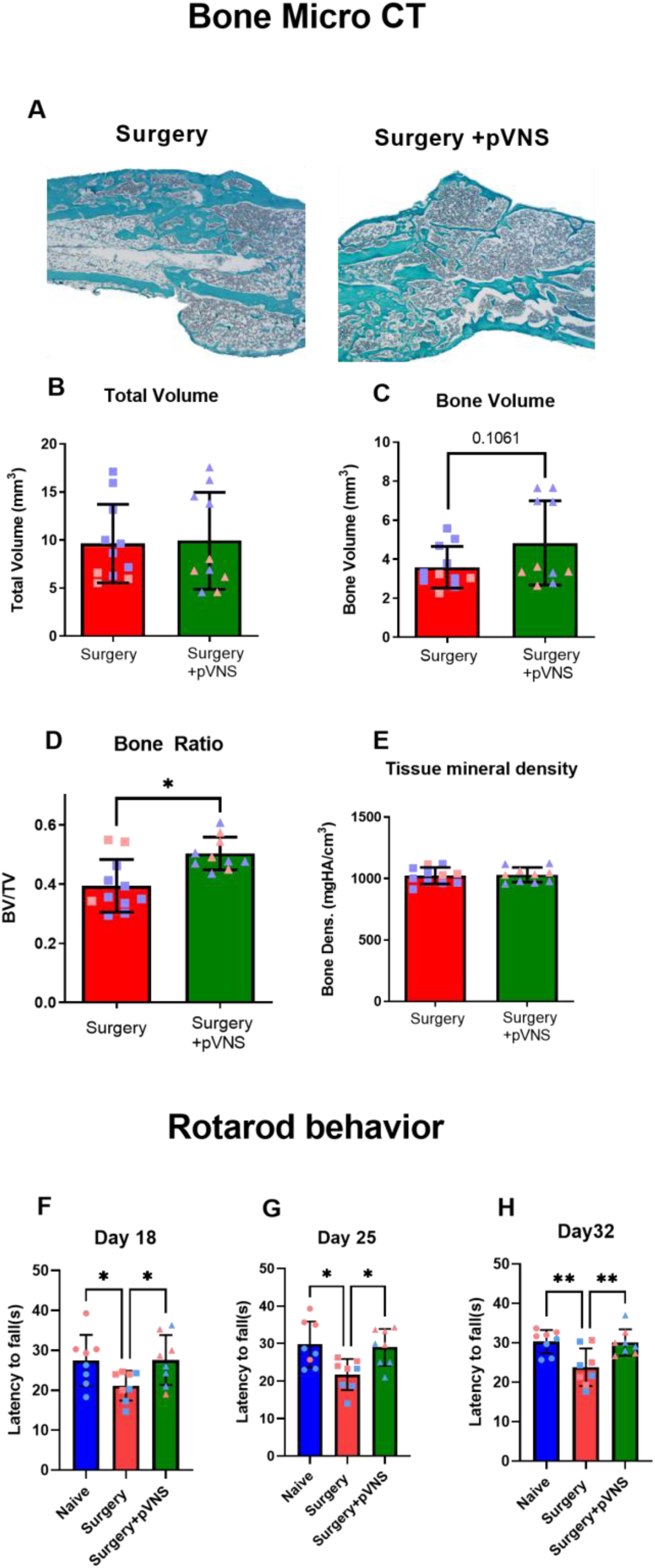
Effects of pVNS on post-surgery bone healing and rotarod behavior. (A) represented bone image for surgery and surgery+pVNS. Quantification suggested pVNS significantly improve bone ratio (D) compared to surgery only group. Surgery +pVNS group has higher trend to increase bone volume but non-significant. The total volume(B) and Tissue mineral density (E) remain no significant difference. n=4-5/sex/group, analyzed by Unpaired t-test. Improvement of rotarod behavior was found in surgery+ VNS group compared to surgery only group on 18 days(F), 25 days(G), and 32 days(H) postsurgery. Analyzed by one-way ANOVA with Tukey posttest. *, **p < 0.05, 0.01 Surgery vs. Sham or Surgery vs Surgery+VNS. Both sex was included in each group. N=5-6 in each sex/group.

### 3.5 Repetitive pVNS rescue satellite glial cell activation in DRG

Glial cells activation in DRG is reported in acute to chronic pain transition [12; 27]. Here we examined Iba-1 and GFAP expression in DRG at 22 days after surgery, when the rescuing effect of pVNS was observed. Notably, no difference in Iba-1 expression and morphology was observed in either group after surgery (1way-Anova with Tukey posttest, p=0.9723, MD=−0.4911, 95% CI [−6.284 to 5.301]) (Fig 5. B). However, GFAP immunoreactivity was found significantly increased in the surgical group compared to naïve group (p=0.0006, MD=12.16, 95% CI [18.93 to 5.384]). Interestingly, repetitive pVNS reversed GFAP elevation after surgery (p=<0.0001, MD=−13.31, 95% CI [−7.037 to −19.58]) (Fig 5. B, G). In the DRG we also observed a significant increase in Monocyte Chemoattractant Protein-1 (MCP-1/CCL2) (p=0.0031, MD=19.24, 95% CI [31.63 to 6.848]), a chemokine involved with persistence pain, which was also reduced by pVNS treatment (p=0.0072, MD=−17.2, 95%CI [−4.812 to −29.60]) (Fig. 5 H). Both MCP-1 and GFAP expression in DRG are highly colocalized as shown by high Pearson Correlation Coefficient (0.73 and 0.783 in Surgery and Surgery+VNS, respectively) (Fig. 5 I). Together, these data highlight a new role for pVNS in regulating satellite cells activation at a key timepoint of conversion from acute to longer-lasting pain in this model.

**Fig 5.**
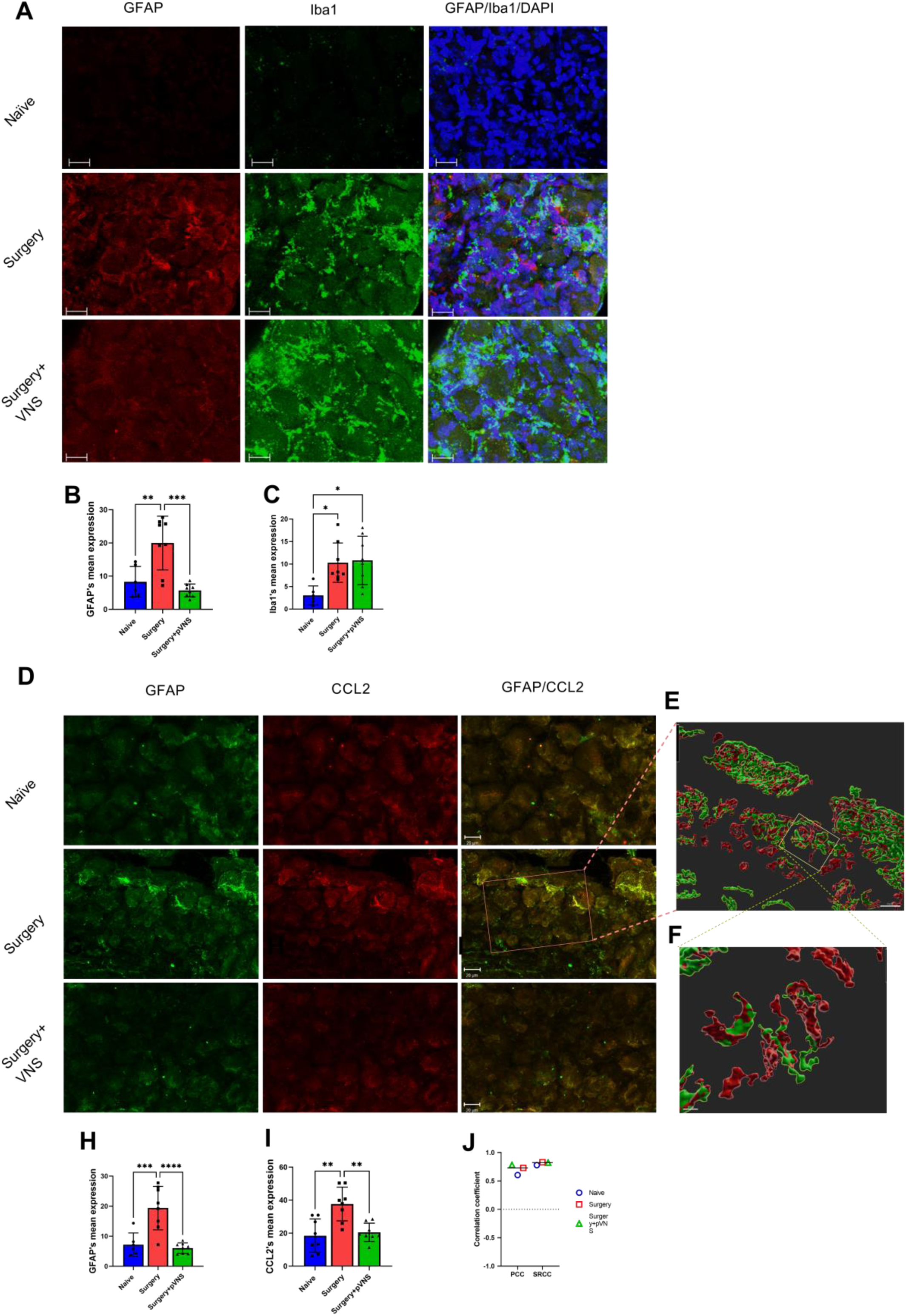
pVNS reduces GFAP(glial satellite cells) but not iba1(microglia) expression in DRG and CCL2 was highly co-localized with GFAP. (A) Surgery increase GFAP and Iba1 expression on mice DRG was observed on 21 days post surgery. pVNS reduced GFAP expression but not iba1 expression on DRG. Scale bar =20μm. Quantification of GFAP(B) and iba1(C) expression on DRG. (D) Surgery induced both GFAP and CCL2 expression in L2-S1 DRG on 21 days post-surgery and both markers were reduced by multiple pVNS treatments. Scale bar =20μm. (E) 3D surface reconstruction image of GAFP and CCL2 with zoom in(F). Scale bar =20(E), 10(F) μm. Quantification of GFAP(G) and CCL2(H) expression on DRG. (I) The Pearson and Spearman Rank Correlation Coefficient score for Naïve, Surgery and Surgery+VNS groups. N=8-10, one-way ANOVA followed by Tukey post test. (*p < 0.05, **p < 0.01, ***p < 0.001, ****p <0.0001).

## Discussion

This study aimed at testing the efficacy of a novel bioelectronic approach using pVNS on postoperative pain. We found that orthopedic surgery contributed to persistent pain and that repetitive pVNS treatments were able to restore mechanical allodynia and reduce MCP-1 and GFAP expression in DRG 22 days after surgery (when pVNS improved behavioral outcomes).

Using this orthopedic surgery model we have observed allodynia behavior in both male and female adult mice for up to 73 days following the procedure. Muwanga et. al. with a similar model for studying orthopedic surgery-induced pain showed more than 5 weeks mechanical allodynia [38]. Many previous groups have reported that unilateral injury model induced contralateral pain-like behavior including animal and clinical models [2; 26; 45; 51]. Enax-Krumova et. al. also reported that patient with unilateral painful or painless neuropathy change contralateral pain sensory [14]. In this study, we have similar result of bilateral side allodynia following unilateral tibial fracture surgery.

Treatment options for postoperative pain remain limited. NSAIDs and opioids are the most common and effective therapies to manage post-surgery pain. Both NSAIDs and opioids have established side effects. NSAIDs have been reported to delay bone formation and decrease bone healing following surgery or fracture [1; 60]. NSAIDs have also been associated with an increase in the risk of falling [64]. Opioids are also known to inhibit callus formation and increase risk of falls, including secondary fractures, during the postoperative recovery [7; 43]. In addition, studies have shown that motor impairments are a common side effect of opioid treatment, which significantly increase falls and fracture risk with odd ratios 1.51 [45]. Lastly, opioid addiction and the ongoing epidemic are well-acknowledged phenomenon that urge development of safe and effective treatments for pain.

Here we have applied a bioelectronic approach leveraging electrical stimulation of the vagus nerve (pVNS) as a translatable intervention to treat postoperative pain. pVNS in this model was effective in preventing prolonged mechanical allodia without significant side effects. In fact, multiple pVNS showed a slight improvement on the rotarod test (Fig 4). The vagus nerve provides innervations to the bone marrow and the parasympathetic system nuclei via cholinergic fibers to bone local cholinergic receptors [6; 50]. To date few studies have addressed the impact of VNS on bone. A multicenter clinical study found that VNS increased femoral and lumbar bone mass density in patients with refractory seizure [54]. To our knowledge, no other preclinical study has analyzed the impact of VNS on bone formation, and this warrants more research given the growing use of VNS to treat multiple immune conditions. Notably, we observed that pVNS increased both bone ratio and mass in mice postoperative model. Research from Tamimi *et. al.* hypothesize that VNS triggers the thyroid gland to decrease T3 and subsequently decrease osteoclast activation [54]. In addition, studies reported that acetylcholine play a regulatory role in both the proliferation and differentiation of osteoblasts [46]. In our research, pVNS dramatically decreases CCL2 in the peripheral nervous system. Earlier studies highlighted an important role for CCL2 to involve bone metabolism [48]. Loss of CCL2 expression in mice leads to a significant increase in bone mass and serum collagen type I fragment. Rat models also demonstrated that CCL2 can be an early predictor of bone loss [22]. In vitro, CCL2 deficiency resulted in reduced numbers and activity of osteoclast [53]. Given the effects we observed on CCL2 in DRGs following pVNS treatment, it is possible that modulation of this target in other sites could account for some of these healing benefits after stimulation.

Vagal stimulation has well-established effects on cholinergic signaling, as the vagus is a key regulator of the cholinergic anti-inflammatory pathway [4; 57]. The identification of the “inflammatory reflex” provided the first description of a neural circuit capable of providing information in real-time to the brain about the body’s inflammatory status, allowing rapid neural regulatory responses via signaling from the vagus nerve and cholinergic singling [40; 56]. Previous work from our group indicated that right pVNS induced bilateral c-Fos activation in the DMX and NTS [24]. Here we have applied the same stimulation titrated to avoid serious bradycardia adverse events and validate the activation of this circuit using FosTRAP reporter system and 20 Hz stimulation frequency. We then applied this protocol and weekly stimulations to ascertain the effects on persistent pain behavior after surgery. Here, we found that weekly pVNS resolved postoperative allodynia on day 22 after surgery. Notably, it took three weeks and four pVNS treatments to show benefit on mechanical allodynia on both paws. The potential key factor(s) to explain this result include: ***(1)*** the necessity of repetitive VNS to modulate specific factors to ultimately improve pain behavior; ***(2)*** a necessary timeframe for bone healing and pVNS impacting these endpoints, thus improving pain recovery; ***(3)*** a detection and/or sensitivity limit of monofilament with up-down method to examine pain-like behavior. Neuromodulation holds many promises, and although some articles discuss the effect of VNS for headache or/and migraine, very few studies have applied VNS on peripheral insult induced chronic pain [32; 37]. Thus, there is an urgent need to understand and optimize the parameters for VNS to achieve optimal pain management and sustained analgesic effects.

Glia play critical roles in chronic pain. In the central nervous system, microglia in the spinal cord dorsal horn correspond with several pain related marker such as Caspase-6, TNF and IL-1β [18; 33]. Astrocytes perform important functions for neuron post-injury response and enhance chronic pain. For example, GFAP upregulation was reported to be highly correlated with pain hypersensitivity after a variety of intervention, including injection of complete Freund adjuvant (CFA)[42], cancer[47], chronic opioid exposure[31], and neuron injury like chronic construction injury (CCI) or spinal nerve ligation (SNL) in rodents[16; 17; 39]. Inhibition of GFAP expression reduced neuropathic pain behavior after nerve injury [58]. However, few reports analyze the relationship of GFAP, a Satellite glial cells (SGCs) marker, in DRG and persistence pain. Kim *et al.* found that GFAP expression in DRG was increased after SNL procedure and that GFAP knockout mice had shorter and less intense pain behavior compared to wild types [34]. Here, we confirmed that the GFAP expression in DRG was elevated after tibial surgery compared to naïve group indicating that SGCs may contribute to the persistence allodynia in our model. In addition, multiple pVNS were able to reduce SGCs activation. Using 3D reconstruction, we further illustrate that CCL2 expression is highly co-localized with GFAP with 0.73 and 0.78 Pearson correlation coefficient in surgery and surgery+VNS group, respectively. Different injury, including chemotherapy and nerve injury, have been reported to induce the CCL2 marker increase in the peripheral system, particularly in DRG [13; 25; 61]. Indeed, CCL2 expression was increased after our orthopedic surgery model and multiple pVNS treatments reduced its expression (Fig 5). The expression of CCL2 is correlated with allodynia behavior in our experiment. Thacker *et. al.* validated that CCL2 induces rodent’s pain states, whereas CCL2 neutralizing antibody reverses mechanical allodynia[55]. This further confirms how the close glia and inflammatory modular play an important role in persistent pain.

In conclusion, our study used a clinically relevant postoperative mice model to examine a minimally invasive, ultrasound-guided pVNS effect on post-surgery pain behavior. We demonstrated that weekly pVNS resolved postoperative-induced bilateral allodynia after 22 days of surgery. We also showed that pVNS have benefit on bone healing and improve the mice motor behavior. In DRG, the expression of the GFAP, a SGCs marker, and CCL2, a pro-inflammatory marker, was increased after surgery but wasreduced with pVNS treatments. The colocalization analysis between GFAP and CCL2 provides evidence of SGCs and CCL2 interaction in DRG. Our findings support the analgesic benefits of pVNS with minimal side effects on chronic pain and urge the importance of SGSc effect on pain management.

## Acknowledgements

This project was supported by grants P01AT009968 of the National Center for Complementary and Integrative Health (NCCIH) and R01-AG057525 of the National Institute on Aging (NIA), the National Institutes of Health (NIH). The sponsors had no role in the design and conduct of the study; collection, management, analysis, and interpretation of the data; preparation, review, or approval of the manuscript; and decision to submit the manuscript for publication. The content is solely the responsibility of the authors and does not necessarily represent the views of the NIH.

